# AAV-Mediated Ribozyme Targeting of HSV-1 Latency-Associated Transcript RNAs Reveals Distinct Functions of the LAT Intron and Exon *In Vivo*

**DOI:** 10.64898/2026.10.13.758874

**Authors:** Mason A. Shipley, Jasbir Singh, David C. Bloom, Donna M. Neumann

**Author notes:** **Co-Corresponding Authors:** David C. Bloom, Ph.D., Department of Molecular Genetics and Microbiology, University of Florida, Gainesville, Florida,; Donna M. Neumann, Ph.D., Department of Ophthalmology and Visual Sciences, University of Wisconsin-Madison, Madison, Wisconsin.

## Abstract

Herpes Simplex Virus 1 (HSV-1) establishes lifelong latency in sensory neurons of the trigeminal ganglia (TG), from which periodic reactivation causes recurrent disease. The latency-associated transcript (LAT) has been implicated in latency establishment, neuronal survival, and reactivation; however, most functional studies have relied on viral deletion mutants that alter genomic architecture and disrupt overlapping noncoding RNAs, making transcript-specific functions difficult to define. To investigate LAT RNA function independent of viral genome modification, we used recombinant adeno-associated virus (rAAV) vectors to deliver hammerhead ribozymes targeting either the LAT 5′ exon or the stable LAT intron in a rabbit model of HSV-1 infection. We found that knocking down the LAT intron prior to HSV-1 infection significantly increased acute disease morbidity and reduced latent viral genome loads in the TG, whereas depletion of the LAT 5′ exon had no detectable effect on latency establishment. LAT intron targeting was also associated with reduced abundance of both the intron and exon LAT transcripts during latency. In contrast, post-establishment depletion of either the LAT 5’ exon or intron significantly reduced reactivation-associated virus shedding following induced reactivation. These findings provide direct evidence that LAT-derived RNAs contribute to multiple stages of the HSV-1 life cycle. Furthermore, the differential effects of LAT exon and intron depletion suggest that functions required for efficient latency establishment localize more closely to the LAT intron, whereas both LAT exon and intron transcripts contribute to efficient reactivation. More broadly, this study demonstrates the utility of transcript-targeting approaches for dissecting the functions of complex herpesvirus noncoding RNA loci.

**Importance:** The HSV-1 latency-associated transcript (LAT) is the only abundantly expressed viral transcript during latency and has been linked to latency establishment, neuronal survival, and reactivation. However, defining the functions of individual LAT-derived RNAs has been challenging because most studies have relied on viral deletion mutants that disrupt overlapping regulatory elements. Using an RNA-targeting approach that preserves viral genome integrity, we demonstrate that the LAT intron promotes efficient establishment of latent infection, whereas both LAT exon and intron RNAs contribute to reactivation-associated virus shedding. These findings provide direct evidence that the LAT RNAs themselves, rather than the underlying DNA elements, regulate key stages of the HSV-1 life cycle.

## Introduction

Herpes Simplex Virus 1 (HSV-1) is a ubiquitous human pathogen infecting ∼70% of adults worldwide (1, 2). Primary lytic infection establishes lifelong latency within sensory neurons, from which periodic reactivation can cause recurrent disease ranging from herpes labialis to keratitis, corneal blindness, and life-threatening encephalitis (3–6). During latency, the viral genome is transcriptionally silenced except for the Latency-Associated Transcript (LAT) locus. The primary LAT primary transcript consists of an ∼6.3 kb unstable exon and a highly stable 1.5 and 2.0 kb intron lariats (7–9). The LAT locus also encodes multiple small noncoding RNAs (sncRNAs) and microRNAs (miRNAs) expressed across different stages of the viral life cycle (10–12). Extensive genetic studies over past decades have implicated the LAT in neuronal survival, latency establishment, genome maintenance, and reactivation (7, 13–24). However, because the LAT encodes multiple overlapping noncoding elements, the precise molecular mechanisms by which the LAT RNA itself contributes to these phenotypes remain poorly understood.

Much of our understanding of LAT functions stems from viral deletion mutants, but this strategy carries inherent limitations. Deletions within the LAT genomic region frequently disrupt adjacent or overlapping regulatory elements and noncoding RNAs, confounding whether observed phenotypes result from loss of the LAT RNA itself or collateral genetic disruption. For example, the LAT-null virus 17N/H exhibits delayed reactivation and reduced plaque size, but its deletion encompasses the LAT 5′ exon, stable intron, adjacent sncRNAs 1 and 2, and miR-H15, while also generating aberrant truncated transcripts and a 29 kDa polypeptide (10, 25). Similarly, the promoter deletion mutant 17ΔPst markedly reduces LAT expression but simultaneously abolishes expression of multiple key miRNAs, including miR-H1, miR-H2, H4-3p, H4-5p, H5, and H6 (19, 20, 26). Consequently, attributing specific functions strictly to primary LAT RNA transcripts versus neighboring genomic or transcriptomic changes remains challenging.

Despite these caveats, deletion studies have provided preliminary evidence that distinct subregions of the LAT execute specialized roles. The LAT 5′ exon region is predominantly implicated in promoting efficient reactivation (13, 15, 26–29). In contrast, the stable LAT intron has been associated with anti-apoptotic activity, cell viability, and latency maintenance (21, 30, 31). Neuronal apoptosis *in vivo* is significantly elevated in mice infected with intron-deletion mutants, whereas 5′ exon deletion mutants retain anti-apoptotic activity (21, 23, 30, 32). Conversely, deletion mutants targeting parts of the intron (e.g., 17Δ480, 17Δ489, 17Δ969) still reactivate following epinephrine induction, indicating that the stable intron is not strictly required for reactivation (27) and recently, the LAT intron region was shown to maintain the latent genome through physical interactions with transmembrane protein TMEM43 (31), suggesting distinct functional roles for the LAT 5’ exon and LAT intron.

Post-transcriptional targeting of components of the LAT transcript can directly alter LAT expression levels without modifying the underlying viral genome, providing a crucial complementary approach to dissect LAT function. We previously established an *in vivo* transcript-depletion strategy using recombinant adeno-associated virus (rAAV8) vectors expressing catalytic hammerhead ribozymes (29, 33). Ocular administration of rAAV8 transduces over 75% of trigeminal ganglion (TG) neurons without inducing ocular pathology or unintended reactivation (33). We previously showed that delivering a ribozyme that targeted the LAT 5′ exon in latently infected rabbits significantly reduced 5′ exon expression without altering viral genome copy number or LAT intron expression, and led to a marked reduction in reactivation (29), providing a valuable tool to assess the impacts of targeting components of the LAT transcript for depletion. Importantly, rAAV vectors can be delivered either prior to HSV-1 infection (to evaluate acute infection and latency establishment) or after latency is established (to evaluate maintenance and reactivation), thereby allowing for the temporal dissection of LAT RNA function across the infection cycle.

In this study, we developed and characterized an AAV-delivered hammerhead ribozyme targeting the stable LAT intron (rAAV8-Rz992) and compared its activity to our exon-targeting ribozyme (rAAV8-Rz235), individually and in combination. We evaluated their ability to degrade LAT transcripts and assessed the consequences of LAT intron versus LAT exon depletion during acute infection, latency establishment, and reactivation *in vivo*. Based on deletion mutant studies, we hypothesized that targeting the LAT intron for depletion prior to infection would impair latency establishment. In fact, pre-infection depletion of the LAT intron significantly increased acute infection morbidity, reduced latent genome loads in the ganglia, and resulted in a concurrent decline in both the LAT intron and LAT 5’exon RNAs. Unexpectedly, post-establishment of latency, targeting of the LAT intron significantly reduced infectious virus shedding during reactivation, like that observed following post-infection treatment with the rAAV8-Rz235. These findings demonstrate that the LAT intron directly contributes to multiple and distinct stages of the HSV-1 life cycle and highlights the utility of transcript-targeting strategies for functional dissection of complex viral loci.

## Results

### Validation of the LAT Intron Targeting Ribozyme (rAAV8-Rz992)

Hammerhead ribozymes (Rzs) are small catalytic RNA molecules featuring a conserved catalytic core flanked by customizable hybridizing arms that mediate sequence-specific substrate cleavage. To target the stable LAT intron of HSV-1, potential hammerhead cleavage triplets (NUH motifs) within the intron sequence were identified, and complementary antisense flanking arms were designed to hybridize to the region surrounding the selected target site (position 992). The resulting sequence was cloned into an AAV2-based transfer plasmid encoding a green fluorescent protein (GFP) reporter driven by a CMV promoter to facilitate assessment of transduction efficiency (**Fig. 1**).

**Figure 1:**
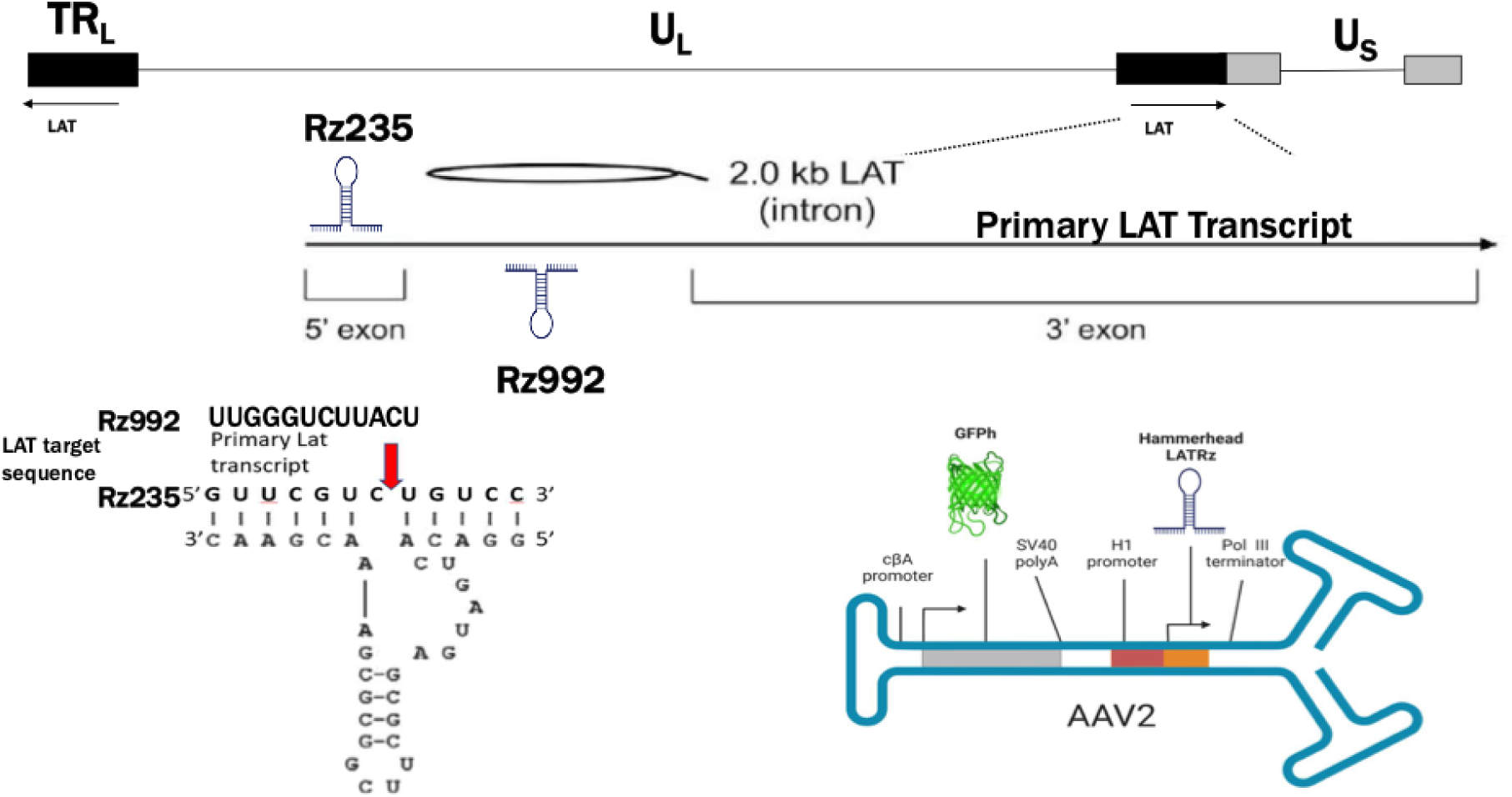
The HSV-1 genome showing the regions encoding the HSV-1 LAT, and design of a ribozyme targeting the LAT intron portion of the LAT RNA. The LAT locus (located in both copies of the R_L_ region of the genome) is expanded to show the 8.5 kb primary LAT, the 5’ and 3’ exons and the stable 2.0 kb intron. The 2.0 kb intron of the HSV-1 LAT was analyzed for hammerhead cleavage sequences. Antisense flanking regions were designed around the conserved catalytic domain to hybridize with the region surrounding the selected triplet. MFOLD analysis predicted base pairing in stem II (arrow), but not in the flanking regions, so this rules out intramolecular base pairing within the ribozyme interfering with binding of the ribozyme efficiently to its target. The ribozyme coding sequences were cloned into an AAV2 vector backbone behind an h1 promoter. This AAV plasmid was then packaged into a pseudotyped AAV8 vector as described in the materials and methods.

To evaluate target knockdown efficiency *in vitro*, rabbit skin (RS) cells were co-transfected with the LAT intron-targeting ribozyme construct (LAT Rz992) and a LAT expression plasmid. Relative LAT intron transcript levels were quantified by reverse transcription quantitative real-time PCR (RT-qPCR) using TaqMan gene expression assays and normalized to the host housekeeping gene *GAPDH*. Co-transfection with a non-targeting control plasmid encoding an anti-rhodopsin ribozyme (RzRho) served as a negative control. Co-expression of LAT Rz992 resulted in a 58% reduction in LAT intron RNA levels compared to cells transfected with the LAT expression plasmid alone.

### LAT 5’ Exon and LAT Intron Transcripts Promote Reactivation

To evaluate the functional consequence of targeted LAT transcript degradation on reactivation *in vivo*, latently infected rabbits were administered rAAV8 vectors expressing the control anti-rhodopsin ribozyme (rAAV8-RzRho), the LAT exon-targeting ribozyme (rAAV8-Rz235), the LAT intron-targeting ribozyme (rAAV8-Rz992), or a combination of both (rAAV8-Rz235 + rAAV8-Rz992). Following vector trafficking to the trigeminal ganglia (TG), reactivation was induced using transcorneal iontophoresis of epinephrine (TCIE), and ocular shedding of infectious virus was monitored daily via swab collection over a 10-day period (26, 34, 35), as outlined in the experimental timeline (**Fig. 2A**). Infectious virus was recovered from 5 of 6 rabbits (83.3%) in the RzRho control group, compared to 3 of 6 (50.0%) in the Rz235 group, 4 of 5 (80.0%) in the Rz992 group, and 5 of 6 (83.3%) in the combination treatment group (**Fig. 3**). Although the overall proportion of animals that reactivated did not differ significantly between groups, all ribozyme treatment groups demonstrated a significant reduction in the frequency of swabs positive for infectious virus. Positive ocular swabs were detected in 25.8% of collections from RzRho control animals, compared to 6.3% in the Rz235 group (p < 0.0005), 15.0% in the Rz992 group (p < 0.05), and 10.0% in the combination group (p < 0.005; **Fig. 3**). Notably, simultaneous targeting of both LAT exon and intron did not further reduce shedding beyond that observed with Rz235 alone. This lack of an apparent additive effect may suggest overlapping functional contributions of the exon and intron transcripts to reactivation or may reflect a threshold beyond which additional reduction in LAT RNA abundance provides limited benefit. Regardless, these findings demonstrate that depletion of either LAT 5’exon or the LAT intron is sufficient to significantly suppress reactivation-associated shedding.

**Figure 2:**
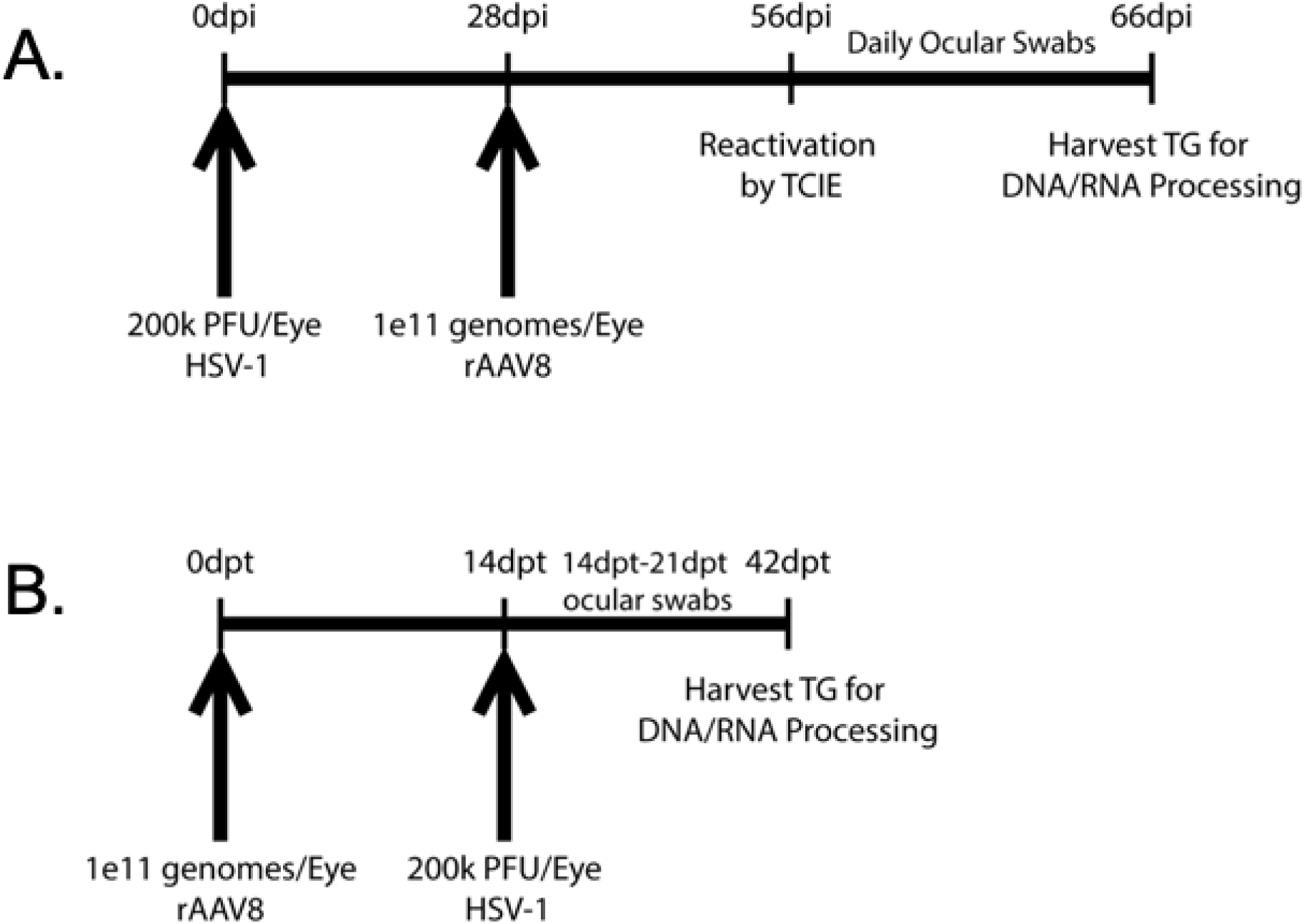
Experimental timelines for either the post-establishment of latency treatment or pretreatment with our ribozyme-containing rAAV vectors. Briefly, rabbits were inoculated at 0 d.p.i. with **A.** 200,000 PFU/eye of HSV-1 strain 17Syn+ (post-establishment experiments). At 28 d.p.i., rabbits were challenged with 1 × 10¹¹ vector genomes (vg)/eye of the indicated rAAV. At 56 d.p.i. rabbits underwent transcorneal iontophoresis of epinephrine (TCIE) and ocular swabs were obtained daily for plaque assays to quantitate viral reactivation. **B.** 1 × 10¹¹ vector genomes (vg)/eye of the indicated rAAV vector were delivered to the surface of the rabbit cornea (pretreatment experiments). At 14 days post vector delivery, rabbits were inoculated with 200,000 PFU/eye of HSV-1 strain 17Syn+. Ocular swabs were obtained for 7 d.p.i and rabbits were monitored for morbidity. At 42 days post-pretreatment, TG were harvested for latent genome load and RNA expression.

**Figure 3:**
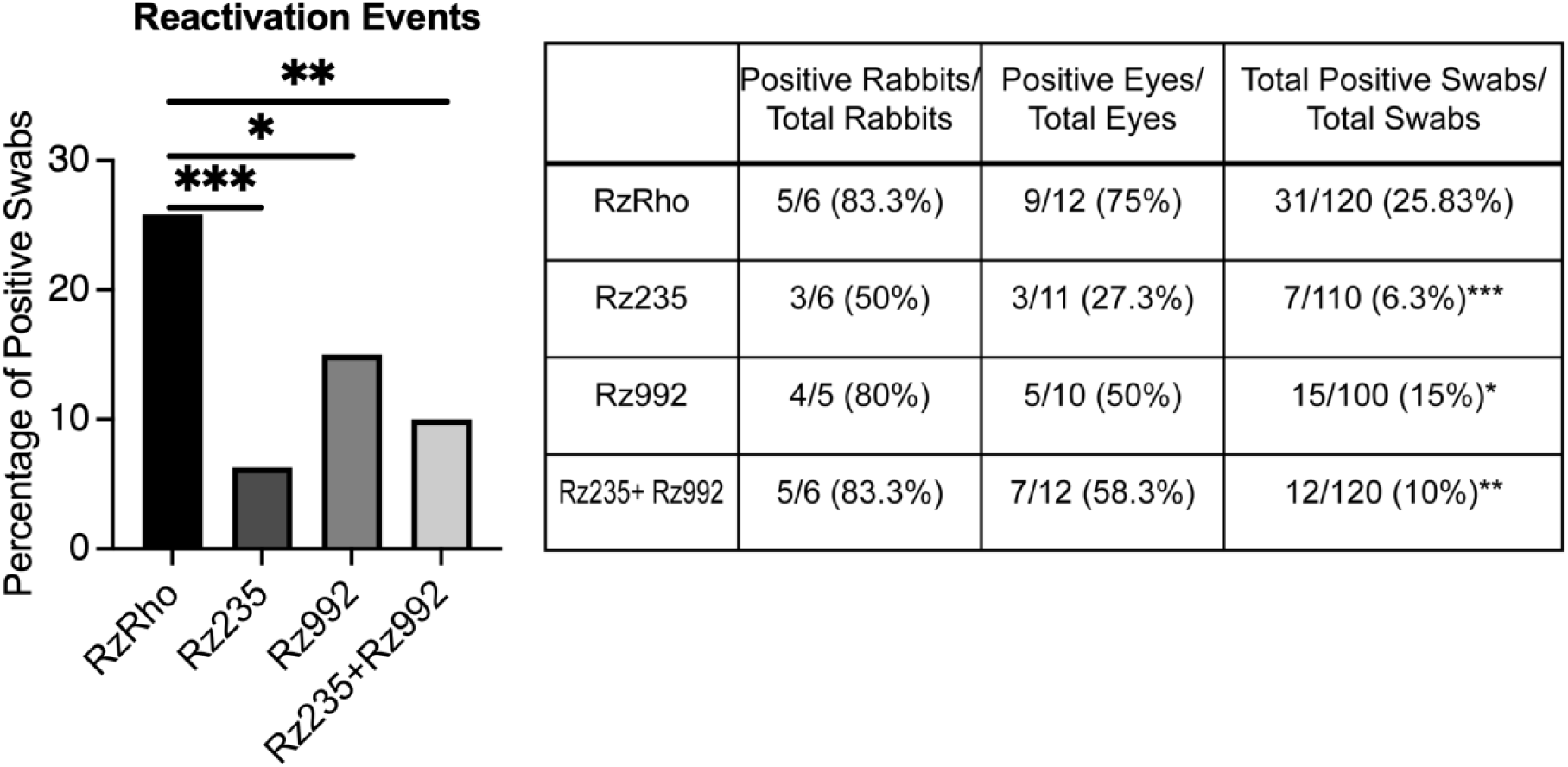
Frequency of HSV-1 reactivation following rAAV treatment and TCIE. Latent rabbits were induced to reactivate by TCIE, and tear-film was collected daily for 10 days post-TCIE. Infectious virus shedding was quantified by plaque assay on vero cells. Compared to the control RzRho, all three rAAV treatment groups exhibited significant reductions in the numbers of virus positive swabs, as determined by chi-square analyses (\**p*<0.05, \*\**p*< 0.005, \*\*\**p*<0.0005).

### LAT Intron Depletion Increases Acute Disease Severity

To determine the contribution of individual LAT subregions during the initial stages of infection and latency establishment, naive rabbits were pre-treated with rAAV8-RzRho (control), rAAV8-Rz235 (exon-targeting), or rAAV8-Rz992 (intron-targeting) prior to corneal inoculation with HSV-1 strain 17Syn+. Following vector delivery and establishment in the TG (**Fig. 2B**), animals were infected, and primary lytic replication was monitored by quantifying ocular viral titers during the first week post-inoculation. Ocular viral loads in the cornea during primary infection did not differ significantly among treatment groups (**Fig. 4A**). However, animals pre-treated with the LAT intron-targeting ribozyme (rAAV8-Rz992) exhibited significantly increased acute morbidity by day 11 post-infection compared to the rAAV8-RzRho control group (*p* < 0.05; Figure 4). Pre-treatment with the exon-targeting ribozyme (rAAV8-Rz235) did not significantly impact morbidity compared to controls (**Fig. 4B**). The increased morbidity observed following LAT intron depletion, despite unchanged ocular viral titers, suggests that the LAT intron contributes to host-virus interactions that influence disease severity independently of productive infection in the cornea. This is consistent with previous studies linking LAT intron to neuronal survival (21, 23).

**Figure 4:**
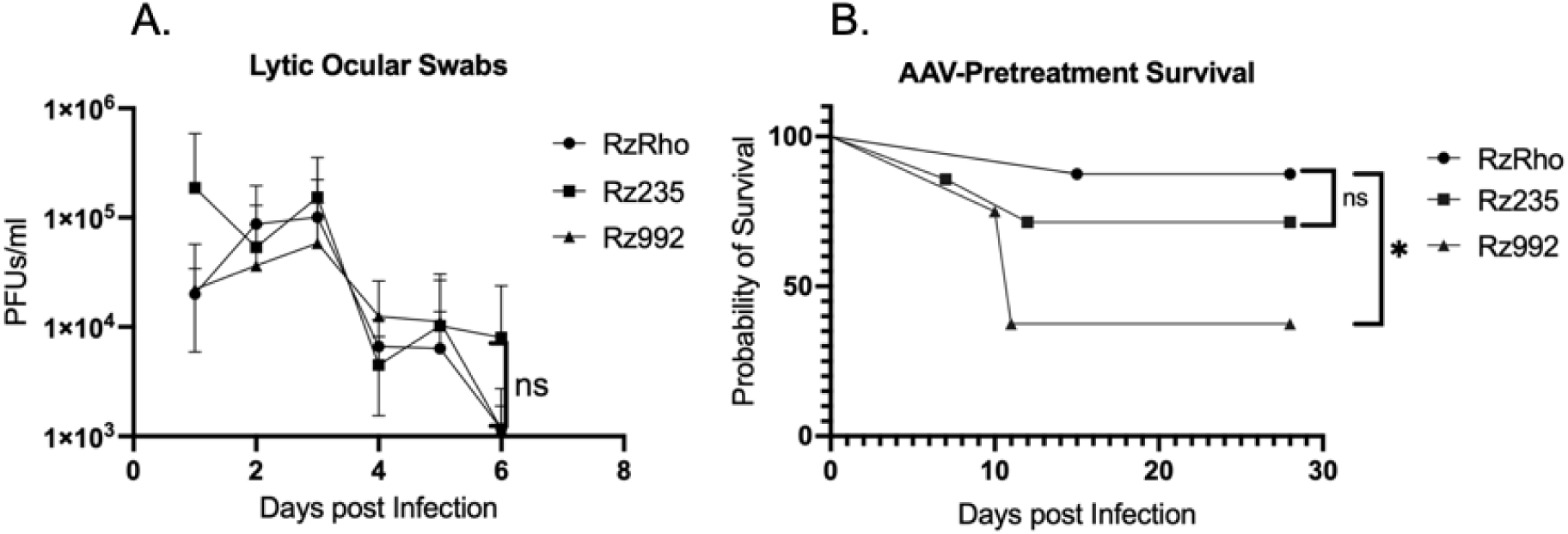
Ocular HSV-1 shedding during acute infection following pretreatment with ribozyme-expressing rAAV vectors; **A**. Rabbits (*n* = 6 per group) were pretreated with the indicated rAAV vectors and subsequently challenged ocularly with HSV-1. Tear-film was collected daily through 7 days post-infection and assayed for infectious virus by plaque assay on vero cells. Data points represent the mean viral titer for each group at each time point ± SD. Statistical analyses were done using two-way analysis of variance. **B**. Kaplan-Meier survival analysis of rabbits (*n* = 6 per group) following ocular HSV-1 challenge after ribozyme pretreatment (\**p*< 0.05).

### LAT Intron Depletion Impairs Latency Establishment

Because LAT intron depletion enhanced acute disease severity, we next assessed the impact of LAT RNA knock-down on the establishment of viral latency. TG tissues were harvested during latent infection, and relative latent viral DNA loads were measured by quantitative PCR targeting the HSV-1 DNA polymerase gene, normalized to host *GAPDH*, and expressed as fold-change relative to the RzRho control group (set to 1.0). Pre-treatment with rAAV8-Rz992 significantly reduced the latent viral genome load in the TG compared to rAAV-RzRho controls (*p* < 0.05; **Fig. 5A**). In contrast, pre-treatment with rAAV8-Rz235 caused no significant change in latent DNA copy number. To determine whether RNA depletion persisted into latency, relative expression levels of both the LAT 5′ exon and the stable intron were quantified by RT-qPCR. Pre-treatment with AAV8-Rz235 reduced 5’exon transcripts ∼2-fold (*p<*0.08), consistent with our previously reported data and we could not detect any 5’ exon expression in the rAAV8-Rz992 treated group, suggesting that depletion of the LAT intron indirectly affects the accumulation of the 5’exon transcript (**Fig. 5B**). In contrast, pre-treatment with rAAV8-Rz235 did not alter LAT intron accumulation while pretreatment with rAAV8-Rz992 resulted in a significant reduction in accumulation of the LAT intron, consistent with LAT intron depletion (*p*<0.05; **Fig. 5B**). Taken together, these data suggest that targeting and degradation of the LAT intron impairs the establishment of viral latency by reducing TG latent DNA load and suppressing LAT transcript expression.

**Figure 5:**
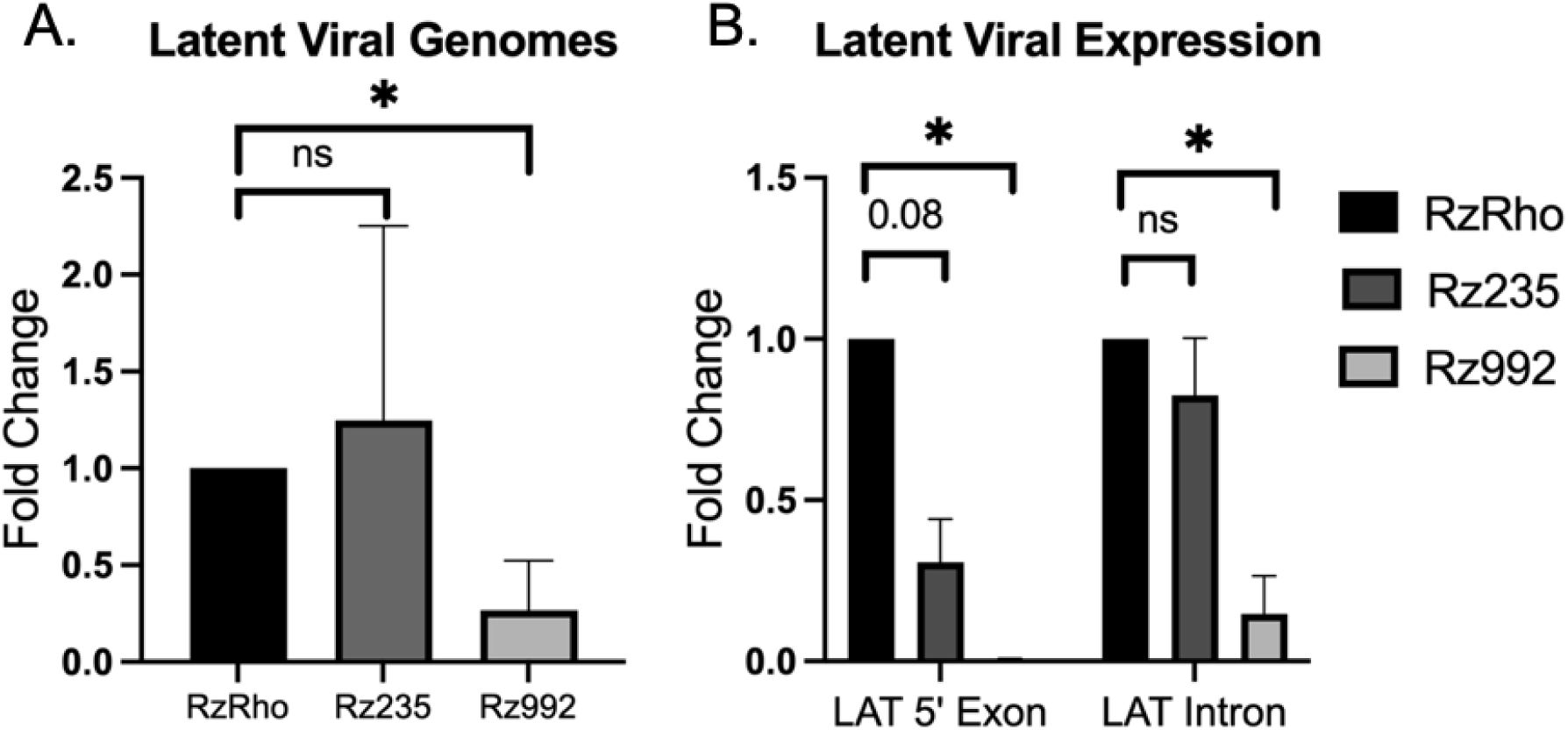
Latent viral genome loads and LAT RNA expression in TG following ribozyme pretreatment. **A.** Relative quantities of HSV-1 DNA Polymerase (DNA Pol) and rabbit Gapdh were determined by qPCR (n=3-4 per group). Viral genome levels were normalized to Gapdh. Graphs are plotted as average fold-change relative to the RzRho control (set to 1) ± SD. Statistical analyses were done using One-Way Analyses of Variance (* p<0.05). **B**. Relative expression of the LAT 5’ exon and stable LAT intron during latency were determined by RT-qPCR for each treatment group (n=3-4 per group). LAT RNA levels were normalized to Gapdh expression. Graphs are plotted as average fold-change relative to the RzRho control (set to 1) ± SD. Statistical analyses were done using One-Way Analyses of Variance (* p<0.05).

## Discussion

The HSV-1 LAT locus encodes a complex collection of overlapping noncoding RNAs, miRNAs, and regulatory elements, making it difficult to distinguish functions attributable to LAT transcripts from those resulting from disruption of the surrounding genomic architecture. To address this limitation, we employed rAAV-delivered hammerhead ribozymes to selectively deplete LAT RNAs without altering the viral genome. Using this approach, we demonstrate that depletion of the stable LAT intron influences both the establishment and reactivation phases of infection. Specifically, pre-infection knock-down of the LAT intron reduced latent viral genome loads within the TG and was associated with decreased LAT transcript abundance during latency. In addition, post-establishment depletion of either the LAT intron or LAT 5′ exon significantly reduced infectious virus shedding following induced reactivation. Together, these findings provide direct evidence that LAT RNAs themselves contribute to multiple stages of the HSV-1 life cycle and highlight the utility of RNA-targeting approaches for dissecting functions within complex viral noncoding RNA loci.

A key finding of this study is that depletion of the LAT intron prior to infection significantly reduced the latent viral genome burden established within the TG. These observations are generally consistent with previous studies implicating the LAT region in neuronal survival, latency establishment, and maintenance of latent genomes (21, 22, 30). However, interpretation of those studies has been complicated by the extensive overlap between LAT transcripts and neighboring regulatory elements. In contrast, the ribozyme strategy used here preserves viral DNA architecture while selectively reducing RNA abundance, providing more direct evidence that LAT-derived RNAs contribute to successful establishment of latency. These findings are particularly interesting in light of recent reports demonstrating physical interactions between the LAT intron, the HSV genome and host proteins involved in latent genome maintenance, including TMEM43 (31). Our results support a model in which LAT intron accumulation promotes efficient establishment or stabilization of the latent viral reservoir within sensory neurons.

The differential effects of knocking down the LAT 5’ exon and intron on latent genome establishment provide additional insight into the functional organization of the LAT locus. While depletion of the LAT intron significantly reduced latent viral genome burden, exon targeting produced no detectable effect on latency establishment despite effectively reducing reactivation-associated shedding. These findings suggest that functions required for efficient establishment of the latent reservoir map more closely to the LAT intron than to the 5′ exon transcript. Previous deletion studies have implicated both regions in latency-associated phenotypes, but interpretation has been complicated by disruption of overlapping transcripts and regulatory elements. By selectively targeting RNA abundance without altering viral DNA architecture, the present study provides additional evidence that latency establishment and reactivation can be functionally distinguished and may depend on distinct LAT-derived RNAs.

An additional and unexpected observation was that targeting the LAT intron prior to infection resulted in a marked reduction in both LAT intron and 5′ exon transcript abundance during latency. Interpretation of this finding is complicated by the concurrent reduction in latent viral genome burden observed following rAAV8-Rz992 treatment. Importantly, although Rz992 was designed to target a sequence within the LAT intron, the target site is also present in the primary unspliced LAT transcript. Consequently, Rz992 is predicted to cleave both the primary LAT transcript and the excised stable intron lariat, making it difficult to distinguish whether the observed reduction in LAT RNAs reflects direct degradation of LAT transcripts, altered processing of the LAT transcriptional unit, or indirect effects resulting from reduced latent genome establishment. Thus, the decrease in 5′ exon-containing LAT transcripts does not necessarily indicate that the intron itself regulates accumulation of these RNAs but may instead result from cleavage of the common precursor transcript before splicing. Alternatively, the reduction in detectable LAT transcripts may simply reflect the presence of fewer latent viral genomes available for transcription. Despite these limitations, the observation that targeting this region reduced both latent genome burden and LAT transcript abundance further supports the conclusion that functions important for efficient latency establishment map within or near the LAT intron region. Moreover, because disruption of this region affects both latent genome establishment and LAT RNA accumulation, the data are consistent with the possibility that critical latency-promoting functions reside within sequences present in the primary LAT transcript and retained in the stable intron.

Unexpectedly, depletion of the LAT intron prior to infection also resulted in increased acute disease morbidity despite no detectable differences in ocular viral titers during primary infection. This dissociation between viral replication at the site of inoculation and disease severity suggests that LAT-dependent functions may influence neuronal survival or host responses independently of productive replication in the cornea. Previous studies have reported increased neuronal apoptosis and enhanced virulence following infection with LAT-deficient viruses (21, 22, 30), supporting the possibility that loss of intron-mediated functions compromises neuronal survival during the transition from lytic infection to latency. Although we did not directly measure apoptosis or neuronal loss, the combination of increased morbidity and reduced latent genome burden is consistent with the long-standing hypothesis that the LAT intron promotes survival of infected neurons during acute infection (21, 23, 30, 32).

We also found that depletion of the LAT intron following latency establishment significantly reduced reactivation-associated shedding. This result was somewhat unexpected given previous reports that several intron deletion mutants retained the capacity to reactivate following epinephrine induction (27). One explanation for this apparent discrepancy is that genomic deletion approaches and transcript depletion approaches interrogate distinct aspects of LAT biology. Deletion mutants alter local genomic architecture and can preserve residual transcript activity while simultaneously affecting neighboring regulatory elements, making transcript-specific conclusions difficult. In contrast, ribozyme-mediated degradation directly reduces RNA abundance while preserving the viral genome. Our findings therefore suggest that the LAT intron contributes quantitatively to efficient reactivation, even if it is not absolutely required for reactivation to occur. Moreover, the observation that intron depletion reduced the frequency of shedding to a degree like exon depletion indicates that both LAT-derived transcripts contribute to optimal reactivation efficiency.

Notably, simultaneous targeting of both LAT RNAs did not further reduce shedding beyond that observed following LAT 5’ exon depletion alone. Although the basis for this remains unclear, the absence of an obvious additive effect may suggest overlapping functional contributions of LAT exon and intron transcripts during reactivation, may be due to decreased delivery of both rAAV8 ribozymes in concurrent treatment, or may reflect a threshold beyond which further reductions in LAT RNA abundance provide little additional effect on shedding frequency. Because the present study was not designed to distinguish between these possibilities, additional experiments examining the extent of transcript depletion achieved by combined ribozyme treatment will be done later. Nevertheless, these data demonstrate that depletion of either LAT RNA species alone is sufficient to significantly suppress reactivation-associated shedding. Collectively, our findings demonstrate that distinct species of LAT transcripts directly contribute to acute pathogenesis, establishment of latent infection, and reactivation of HSV-1. More broadly, these studies establish transcript-targeting approaches as powerful tools for dissecting the functions of complex herpesvirus noncoding RNA loci and provide a framework for defining the specific contributions of individual LAT RNA species to HSV-1 persistence and recurrent disease.

## Methods

### Viruses and Cells

Vero cells (ATCC CCL-81) were cultured in Dulbecco’s modified Eagle’s medium (DMEM; Corning #10–017) supplemented with 10% fetal bovine serum (FBS; Corning #35-011-CV) and 1% antibiotic-antimycotic solution (Gibco #15240062) in a humidified atmosphere with 5% CO_2_ at 37°C. Cells were expanded at regular intervals when the cells were at ∼60% confluency. Original stocks of the wild-type HSV-1 strain, 17*Syn+* (GenBank accession number <u>NC_001806</u> were obtained from the Bloom Lab at the University of Florida. Virus stocks were propagated on Vero cells using a 0.01 MOI. Cultures were supplemented with Eagle’s minimal essential medium with 1% FBS and 1% antibiotic-antimycotic solution at 37°C for 3–4 days and viruses were harvested by centrifugation followed by two freeze thaw cycles. The final supernatants were aliquoted and stored in −80°C for further use. To determine viral titers, Vero cells were infected in triplicate with 10-fold serial dilutions in DMEM (1% FBS and 1% antibiotic-antimycotic solution) for 72 hrs, plaques were stained by crystal violet and counted.

### Target selection and design of synthetic hammerhead ribozymes

The LAT intron region of the HSV-1 genome (NCBI database accession number NC_001806) was analyzed for hammerhead ribozyme cleavage sites. The nucleotide triplets GUC, CUC, and UUC were specifically selected as potential target sites. Hybridizing arms were designed complementary to sequence surrounding the cleavage site. MFOLD software (Zuker, 2003) was used to predict secondary structure of each ribozyme. Ribozymes predicted to have a correctly folded catalytic core and open hybridizing arms were carried forward to *in vitro* testing. Rz992 was selected as the most catalytically active for the LAT intron target.

### Cloning synthetic ribozymes into AAV plasmids

Selected synthetic hammerhead ribozymes were cloned into scAAV2-GFP plasmids for *in vitro* testing and subsequent vector production. In this vector humanized GFP (GFPh) is expressed from the CBA promoter consisting of the chimeric chicken beta actin proximal promoter plus the immediate early enhancer from cytomegalovirus. The self-complementary AAV (scAAV) plasmid contains a modified TR that results in the generation of a self-complementary copy of the transgene during packaging to improve transduction efficiency. The ribozyme sequence was placed downstream of an H1 promoter and terminated by a PolIII terminator sequence. These mini-genes were synthesized by GenScript with SalI restriction endonuclease sites at each end. AAV plasmid containing the AAV2 TRs and the insert were digested with SalI (New England Biolabs) and ligated. Ligations were transformed into SURE electroporation competent cells (Stratagene) and grown in LB media. Plasmids were isolated and purified by CsCl gradient centrifugation. Insertion was confirmed by PCR and Sanger sequencing. The AAV plasmids were pseudo-typed into AAV8 Y733F modified capsids.

### *In vitro* testing of synthetic hammerhead ribozymes

Lipofectamine 2000 (Invitrogen) was used to co-transfect a plasmid encoding synthetic hammerhead ribozymes and a plasmid encoding the target LAT transcript into rabbit skin cells. Each transfection was performed in triplicate in a 24-well plate format. A plasmid expressing an anti-rhodopsin ribozyme was co-transfected in control wells to account for nonspecific effects. Twenty-four hours post-transfection, RNA was isolated using TRIzol (Ambion) and used for subsequent reverse transcription and qPCR analysis as described below.

### Rabbit inoculation with HSV-1 and establishment of latency

New Zealand White rabbits were anesthetized using intramuscular injections of ketamine (30 to 45 mg/kg [body weight]) and xylazine (7.5 to 11.5 mg/kg [body weight]). Using a blunt-tip 27-gauge needle, a 3-by-3 crosshatch pattern was made on the corneal surface. HSV-1 strain 17Syn+ was applied to each eye in a volume of 20 l (200,000 PFU/eye). Rabbits were monitored, and ocular swabs were performed on postinfection days 1-7 to assess ocular viral load during the acute infection.

### Treatment with vectors

Prior to- or following-the establishment of latency (at 28 days p.i.), either rAAV8-RzRho (control) rAAV8-Rz235, rAAV8-Rz992, or a combination of rAAV8-Rz235 and rAAV8-Rz992, was applied to the corneal surface of rabbits following corneal abrasion at an initial inoculum of 10e10 viral genome equivalents/eye for each AAV vector in a total volume of 30ul. All rabbits were monitored for signs of ocular pathology until the termination of the experiment.

### Epinephrine Induced Reactivation

Rabbit corneas were subjected to TCIE using previously reported methods (26, 34, 35). Briefly, latently infected rabbits were anesthetized using IM ketamine/xylazine and subjected TCIE using a 0.015% epinephrine solution in water at 0.8 mA for 8 min per eye. Rabbits were swabbed for 10 consecutive days following TCIE. At the end of the experiment, rabbits were euthanized, and the TG were rapidly removed and either placed in 10% buffered formalin (histology) or placed in RNAlater (Ambion, Inc., Austin, TX) for RNA and DNA extraction.

### Ocular swabs

Rabbit eyes were swabbed with sterile cotton-tipped applicators that were rubbed across the corneal surface and under the eyelids. Each applicator was immediately placed in a tube containing 1 ml of sterile DMEM supplemented with 1% fetal bovine serum and 2% penicillin/streptomycin.

### Ocular viral titer quantification

The tube containing the applicator was placed on a rocker at room temperature for 1 h. A 24-well plate was plated with Vero cells 24 h earlier. The normal growth medium was removed from the cells, and the medium from each tube was then transferred to a single well in the plate. The plate was placed in the incubator at 37°C and 5% CO2. Plates were monitored under the microscope for plaques and/or cytopathic effects for 7 days.

### RT-pCR

Rabbit TG were isolated and placed in RNAlater and stored according to the manufacturer’s specifications. RNA was extracted by removing RNAlater from samples and adding TRIzol reagent (Sigma-Aldrich) to each sample. Briefly, each TG was homogenized in 1.2 ml of TRIzol and, following the addition of 0.2 volume of chloroform, the samples were centrifuged for phase separation. RNA was precipitated from the aqueous phase using 0.7 volume of isopropanol, according to the manufacturer’s directions. Reverse transcription PCR was performed using the Superscript III Platinum One Step RT-PCR kit per the manufacturer’s instructions in triplicate. Rabbit GAPDH Primer, forward GCA CCA CCA ACT GCT TAG C Primer, reverse CCT CCA CAA TGC CGA AGT G Probe CTG GCC AAG GTC ATC. LAT 5’exon and LAT intron primers were previously reported (29).

### qPCR for genome copies/ganglia

Real-time primer/probe sets specific to HSV-1 DNA polymerase were used to determine HSV-1 genome copies per ganglia. Real-time PCR was performed in triplicate. Rabbit GAPDH Primer, forward GCA CCA CCA ACT GCT TAG C Primer, reverse CCT CCA CAA TGC CGA AGT G Probe CTG GCC AAG GTC ATC. HSV-1 DNA polymerase NC_001806 (65880–65953) Primer, forward AGA GGG ACA TCC AGG ACT TTG T Primer, reverse CAG GCG CTT GTT GGT GTA C Probe ACC GCC GAA CTG AGC A. The relative HSV-1 copies for each sample were further normalized to the relative copies of the host control rabbit GAPDH.

## Statistical analyses

Statistical analysis (One Way analyses of Variance, Two-Way Analyses of Variance and chi-squared analyses) was performed with Prism 11.2 for Mac OS.

## Acknowledgements

This work was supported by the grants NIH/NIAID R01AI134807 and R01AI19006 (DMN), NIH/NIAID R01AI048633 (DCB), NIH/NEI 2T32EY027721 (MAS), the Core Grant for Vision Research from the NIH to the University of Wisconsin-Madison (P30EY016665), and an unrestricted grant from Research to Prevent Blindness (Department of Ophthalmology, University of Wisconsin). The funders had no role in study design, data collection and analysis, decision to publish, or preparation of the manuscript.

